# Hemodynamic responses measured with fNIRS scale with transcranial direct current stimulation intensity up to 6 mA

**DOI:** 10.64898/2026.11.27.767075

**Authors:** Jinglin Mo, Yahia Abdalla, Mohamad FallahRad, Mojtaba Belali Koochesfahani, Kyle Donnery, Santiago Osorio Jurado, Marom Bikson

**Author notes:** **Corresponding Author:** Jinglin Mo, **Address**: No. 119 South 4th Ring West Road, Fengtai District, Beijing, China, 100070, **E-mail:**.

## Abstract

**Highlights:** HC-tDCS up to 6 mA produced dose-dependent hemodynamic responses measured by fNIRS.

HbT and HbO responses increased with stimulation intensity from 2 to 6 mA.

Long-separation fNIRS channels and active sham support a cerebral contribution.

At 6 mA, HbO and HbT continued to increase during sustained stimulation.

**Background:** High-capacity (HC) technology enables tolerated transcranial direct current stimulation (tDCS) with intensities up to 6 mA, above the conventional ≤2 mA range. We used functional near-infrared spectroscopy (fNIRS) to characterize the dose dependence of cerebral hemodynamic responses to HC-tDCS across 2-6 mA.

**Methods:** 14 healthy adults participated in a within-subject, sham-controlled pilot study. Each participant underwent 2, 4, and 6 mA HC-tDCS in blinded, randomized order, followed by sham and 0 mA sessions. fNIRS was recorded continuously for 20 min, including 3 min of rest, 2 min of ramp-up, 8 min of stimulation, 2 min of ramp-down, and 5 min post-stimulation. Changes in oxygenated hemoglobin (HbO), deoxygenated hemoglobin (HbR), and total hemoglobin (HbT) were referenced to the resting baseline and corrected using time-matched 0 mA recordings. Dose-response relationships were examined across the three active intensities during the 8-min stimulation period.

**Results:** HbT and HbO responses increased with 2-6 mA HC-tDCS intensity, with dose-response slopes of 0.46 μM/mA for HbT and 0.56 μM/mA for HbO. The intensity-dependent increases in HbT and HbO were observed across all source-detector distances, with the greatest responses at 6 mA, including longer-separation channels (3.0 and 3.35 cm). At 6 mA, HbT and HbO continued to increase throughout the stimulation period (HbT slope: 0.19 μM/min; HbO slope: 0.17 μM/min).

**Conclusions:** HC-tDCS produced intensity-dependent hemodynamic responses across 2-6 mA, with pronounced responses at 6 mA. Furthermore, these findings are consistent with a vascular stimulation mechanism of tDCS.

## Introduction

Transcranial direct current stimulation (tDCS) is a noninvasive neuromodulation technique that applies weak direct current through scalp electrodes to modulate brain function. Increasing stimulation current produces stronger electric fields in the brain [1,2], and cellular studies demonstrate that neuronal polarization increases with electric field strength [3,4]. Most human tDCS studies limit current to 2 mA or less [5]. High Capacity (HC) electrode technology enables stimulation at higher intensities, including 6 mA, with acceptable safety and tolerability [6], providing an opportunity to investigate physiological responses in human trials beyond the conventional tDCS dose range.

Parallel to its established effects on neuronal activity, tDCS may modulate cerebral hemodynamics through both direct vascular stimulation and neurovascular coupling. Direct effects reflect stimulation of perivascular nerves and vascular endothelium, while indirect effects arise from stimulation of neurons or glia that influence blood flow [7]. In vitro and animal studies demonstrate vascular responses to direct current stimulation [8–11], while human perfusion imaging shows changes in regional cerebral blood flow with tDCS [12,13]. Human perfusion studies have also examined dose dependence, with Jamil et al. reporting intensity-dependent perfusion changes across 0.5-2 mA that were related to predicted regional electric field intensity [14]. However, hemodynamic dose-response is inconsistent across studies [15], including when extending stimulation intensity to 4 mA [16–18]. Thus, the consistency and dose dependence of cerebral hemodynamic responses, particularly at higher stimulation intensities, remain unclear.

Functional near-infrared spectroscopy (fNIRS) enables continuous monitoring of cerebral hemodynamic responses during tDCS by measuring changes in oxygenated hemoglobin (HbO), deoxygenated hemoglobin (HbR), and total hemoglobin (HbT). Previous resting-state fNIRS-tDCS studies have reported heterogeneous hemodynamic responses at conventional stimulation intensities [19].

In this study, we used concurrent fNIRS to investigate cerebral hemodynamic responses to HC-tDCS in healthy adults. Our primary objective was to characterize the dose-response across randomized stimulation intensities of 2, 4, and 6 mA. Additional 0 mA and novel skin-active sham stimulation conditions were included as nonrandomized controls.

## Methods

### Study design and participants

This within-subject, sham-controlled pilot study included 14 healthy adults. Before tDCS stimulation, participants completed a brief questionnaire to collect demographic, medical, and lifestyle information, including age, sex, history of mental or physical illness, alcohol and caffeine consumption, nicotine use, and habitual physical activity. Participants were also screened for conditions that could affect the safety or delivery of stimulation, including local skin lesions, irritation, or other factors that could compromise electrode contact. The study was conducted in accordance with procedures approved by the Institutional Review Board of the City College of New York (protocol 2024-0013-CCNY). All participants provided written informed consent before participation.

Each participant completed three active stimulation sessions at 2, 4, and 6 mA in randomized order and was blinded to stimulation intensity. Stimulation order was determined using a pre-generated randomization schedule, with participants assigned sequentially according to enrollment order. An additional nonrandomized sham session and a 0 mA session were conducted after the randomized active sessions (Figure 1A). During each 20 min session, participants remained seated with their eyes open, fixating on a cross displayed on a screen, and were instructed to minimize movement and remain awake. Each session consisted of a 3 min resting baseline, 2 min ramp-up, 8 min stimulation period, 2 min ramp-down, and 5 min post-stimulation phase, with fNIRS signals recorded continuously throughout (Figure 1B). Skin integrity and stimulation tolerability were assessed after each session using skin examinations and post-session questionnaires.

**Figure 1.**
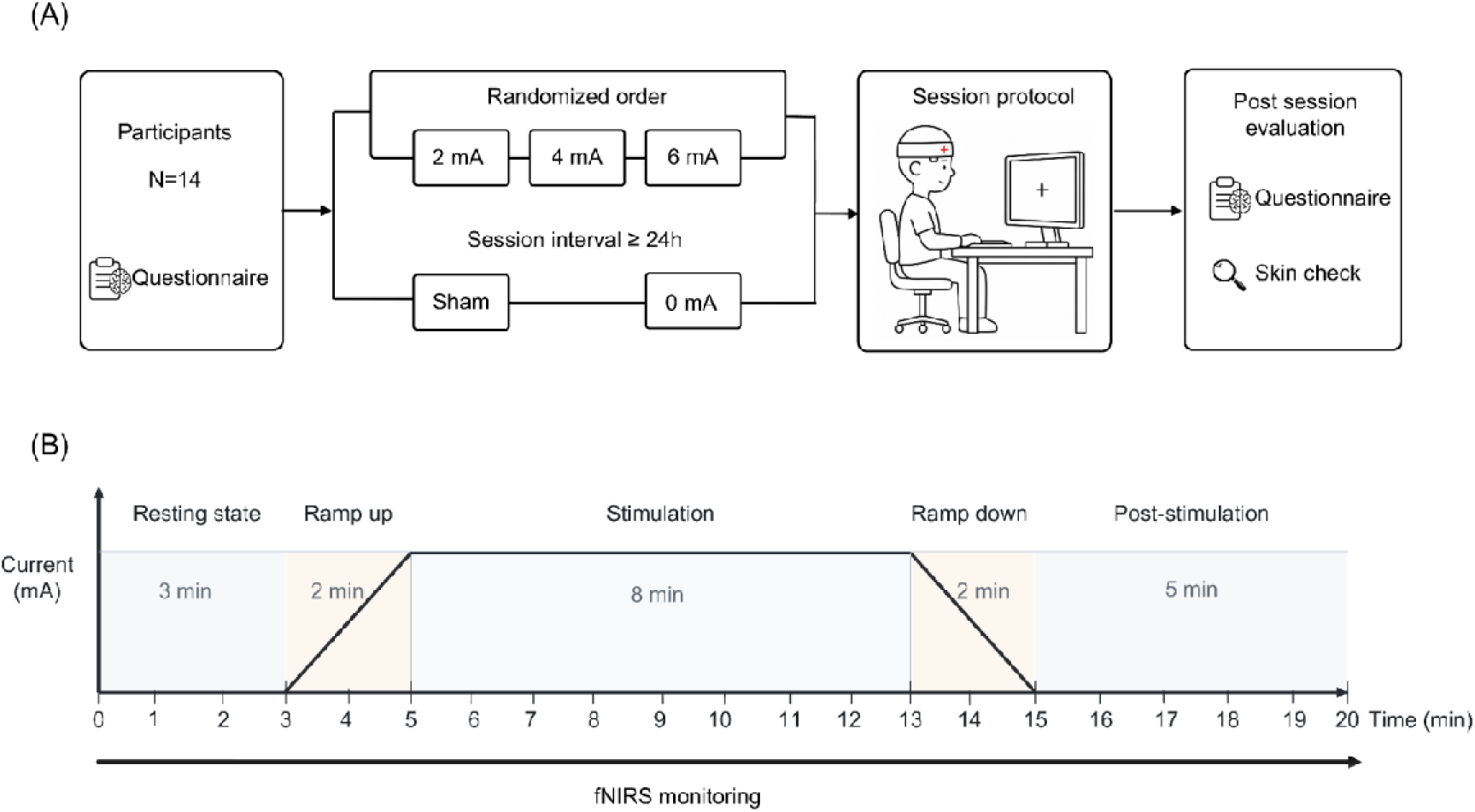
Study design and experimental protocol. (A) Overview of the study design. (B) 20 min stimulation protocol with continuous fNIRS monitoring.

Before stimulation, the forehead skin was cleaned with alcohol wipes, and two HC electrodes were positioned over the left (anode) and right (cathode) frontal regions (Figure 2A). The fNIRS headgear was then placed over the scalp and adjusted around the stimulation electrodes for simultaneous HC-tDCS and fNIRS recording (Figure 2B). Active stimulation was delivered using the previously described HC-tDCS electrode and stimulator system, configured to deliver target currents up to 6 mA. Detailed electrode specifications and corresponding electric-field modeling have been reported [6].

**Figure 2:**
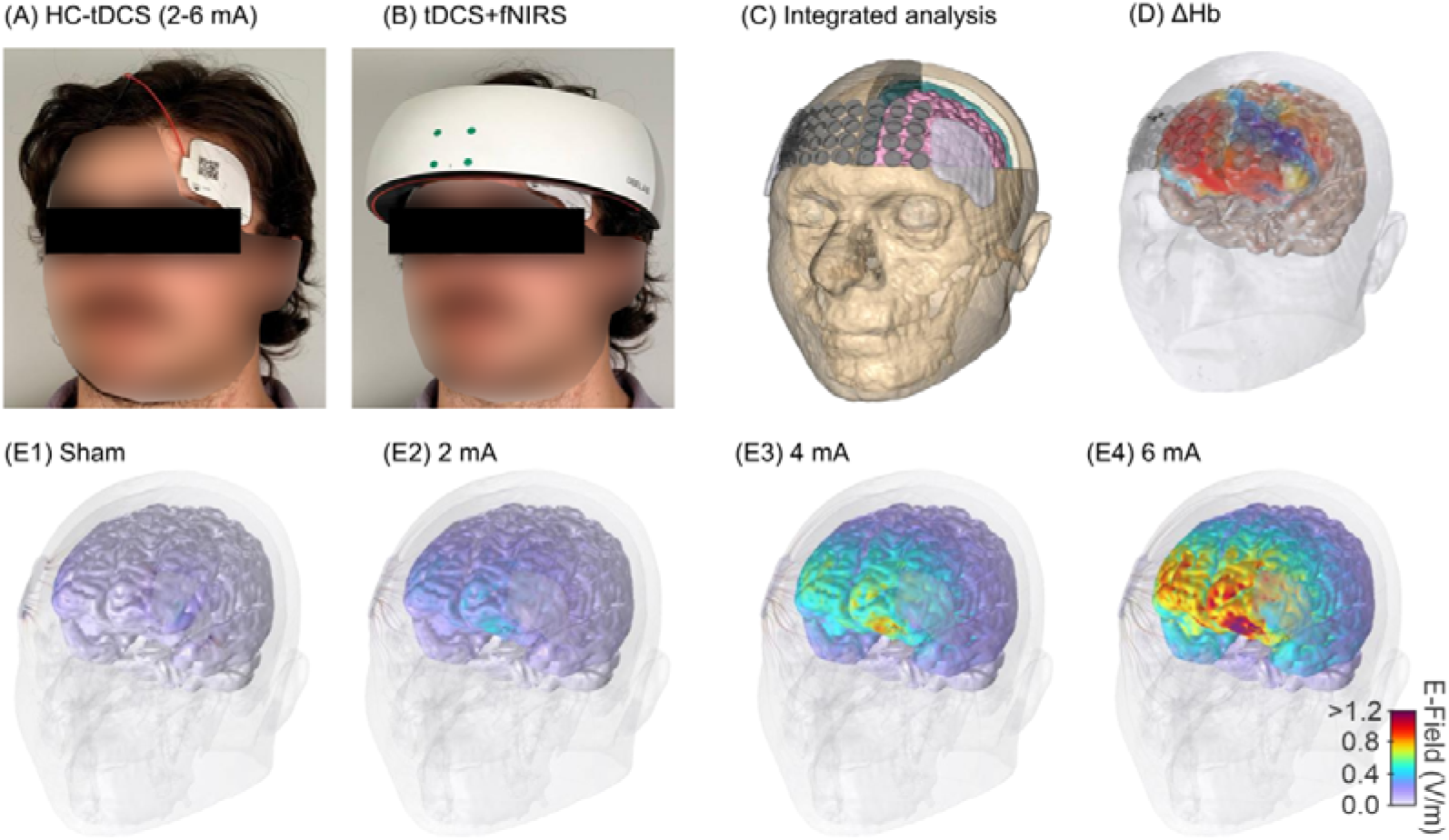
Experimental setup, fNIRS measurements, and electric-field modeling. (A) Placement of HC-tDCS electrodes over the left (anode) and right (cathode) frontal regions. (B) Experimental setup for simultaneous HC-tDCS and fNIRS recording. (C) MRI-derived head model illustrating the spatial arrangement of stimulation electrodes and fNIRS measurement channels. (D) Illustrative spatial map of hemoglobin concentration changes (ΔHb) across fNIRS measurement channels. (E1–E4) Computationally predicted cortical electric-field magnitude distributions for skin-active sham, 2 mA, 4 mA, and 6 mA stimulation, respectively, showing progressively stronger electric fields with increasing stimulation intensity. *The photographs in panels A and B depict one of the authors. The three-dimensional head models shown in panels C, D, and E1-E4 were reconstructed from imaging data acquired from the same author.

In the 0 mA condition, the same electrode assemblies and montage as in the active stimulation conditions were used, but no current was delivered. In the sham condition, the electrode patches had the same external appearance as those used in the active and 0 mA conditions, while being electrically isolated and each containing two closely spaced anodal and cathodal components (Figure S1). This configuration created local current pathways, allowing continuous 1 mA current flow between the paired anodal and cathodal components within each assembly. Unlike conventional tDCS sham protocols, in which current is typically delivered only briefly during ramp-up and ramp-down to produce cutaneous sensations during ramp periods, the present sham condition produces continuous tingling sensation throughout the stimulation period. The close spacing between the anodal and cathodal components was intended to confine most of the current to superficial tissues, thereby reproducing cutaneous sensations while minimizing transcranial current flow and the resulting electric field in the brain.

### fNIRS acquisition and preprocessing

fNIRS data were acquired using the NIRSIT system (OBELAB Inc., Seoul, Republic of Korea) at wavelengths of 780 and 850 nm and a sampling rate of 8.138 Hz. The system comprised 204 source-detector pairs with nominal separations of 1.5, 2.12, 3.0, and 3.35 cm. For illustrative purposes, the spatial arrangement of the stimulation electrodes and fNIRS measurement channels was visualized on a magnetic resonance imaging (MRI)-derived head model from a representative participant, together with an illustrative map of hemoglobin concentration changes (ΔHb) (Figure 2C and D). Raw light-intensity data were exported and preprocessed using the MNE-Python toolbox [20]. Channel quality was assessed using the NIRSIT Analysis Tool (OBELAB Inc.); channels with invalid measurements or optical intensity values below 30 arbitrary units (A.U.) were excluded. Raw intensity signals were converted to optical density, followed by motion correction using Temporal Derivative Distribution Repair (TDDR). The corrected optical-density signals were low-pass filtered at 0.1 Hz. Relative changes in HbO and HbR were then estimated using the modified Beer-Lambert law with age-adjusted differential pathlength factors [21]. HbT was calculated as HbO + HbR.

### Computational modeling and solution method

Electric-field modeling was performed using a finite element head model derived from a high-resolution structural MRI scan (1-mm isotropic voxel size) of one participant, following previously described procedures [6,22]. The MRI was segmented in Simpleware (Synopsys, Inc., Mountain View, CA, USA) into scalp, skull, cerebrospinal fluid, gray matter, white matter, and air. Stimulation electrodes were incorporated according to the experimental montages. A tetrahedral mesh with first-order elements was generated in Simpleware and imported into COMSOL Multiphysics 6.1 (COMSOL, Inc., Burlington, MA, USA). Tissue- and electrode-specific electrical conductivities were assigned, and electric-field distributions were computed using the Electric Currents (ec) interface under stationary, quasi-static conditions. Current-controlled Terminal and Ground boundary conditions were assigned to the corresponding electrodes, accounting for the distinct current pathways in the active and sham configurations. Simulations were performed for sham, 2 mA, 4 mA, and 6 mA stimulation. The final model contained 19,475,834 degrees of freedom. Predicted cortical electric-field distributions are shown in Figure 2E1–E4.

### Statistical analysis

HbO, HbR, and HbT time courses were first averaged across retained channels for each participant and condition. Two forms of the time courses were examined. First, to visualize hemodynamic responses across the randomized active conditions, the 2, 4, and 6 mA time courses were referenced to the mean of their respective initial 3 min resting periods and then averaged across participants. For subsequent quantitative analyses, the 0 mA condition was used to account for temporal changes in the fNIRS signal in the absence of stimulation. Within each participant, the 0 mA time course was subtracted point by point from each sham, 2 mA, 4 mA, and 6 mA time course. Each resulting time course was then referenced to its mean during the initial 3 min resting period. These corrected changes are denoted as ΔHbO, ΔHbR, and ΔHbT throughout the subsequent analyses.

For each condition, mean ΔHbO, ΔHbR, and ΔHbT over the 8 min stimulation plateau was calculated for each participant. The primary dose-response analysis was performed across the randomized 2, 4, and 6 mA conditions by fitting an ordinary least squares regression within each participant. The resulting slopes were tested against zero using one-sample t tests. P values for the HbO, HbR, and HbT dose-response slopes were adjusted across the three hemoglobin measures using the Holm procedure. Sham was not included in the dose-response analysis. As additional comparisons, differences among sham, 2 mA, 4 mA, and 6 mA were assessed using paired t tests. All six pairwise comparisons were evaluated, with P values adjusted across the six comparisons separately within each hemoglobin measure using the Holm procedure. Mean slopes and paired differences are reported with 95% confidence intervals.

The spatial dependence of ΔHbT and ΔHbO was examined at source-detector separations of 1.5, 2.12, 3.0, and 3.35 cm. Mean stimulation period ΔHbT and ΔHbO at each separation were tested against zero. For longer separation signals, channels at 3.0 and 3.35 cm were combined within each participant and condition, and ΔHbO, ΔHbR, and ΔHbT were compared among stimulation conditions using paired t tests. Holm adjustment was applied across the four source-detector separations within each condition and hemoglobin measure, and across the six pairwise condition comparisons within each hemoglobin measure.

Changes in ΔHbT and ΔHbO during sustained stimulation were assessed by fitting a linear slope to each participant’s time course over the 8 min stimulation period (5-13 min). Slopes are expressed in μM/min and were tested against zero for each condition, with P values adjusted using the Holm procedure. Dose dependence of the temporal slopes was examined by a second regression across the randomized 2, 4, and 6 mA conditions within each participant. The resulting dose slopes, expressed in μM/min/mA, were tested against zero and are reported as means with 95% confidence intervals.

All tests were two-sided, with statistical significance defined as P < 0.05. P values are reported after Holm adjustment for analyses in which multiple-comparison correction was applied, as specified above. Analyses were performed in Python using MNE-Python, NumPy, pandas, SciPy, and statsmodels.

## Results

All 14 healthy adults (8 men and 6 women; mean age, 28.5 years; standard deviation [SD], 8.9 years) completed three active stimulation sessions (2, 4, and 6 mA) in randomized order, followed by two nonrandomized control sessions (sham and 0 mA), and were included in the analysis. The low-profile hydrogel HC electrodes allowed fNIRS optode placement over the stimulation regions, although channels directly overlying the optically opaque electrodes were excluded. The mean numbers of retained channels were 87.7 (SD 13.9), 88.6 (15.4), 92.1 (15.0), 85.1 (14.6), and 82.0 (16.1) for sham, 0, 2, 4, and 6 mA, respectively, with no significant difference across conditions (one-way ANOVA, P = 0.466; Table S1).

### Intensity-dependent hemodynamic responses to HC-tDCS

Electric-field modeling in a representative participant showed a linear increase in brain 99th percentile [P99] values from 0.44 V/m at 2 mA to 0.88 V/m at 4 mA and 1.32 V/m at 6 mA (Figure 2E1–E4). We hypothesized that fNIRS-measured hemodynamic responses would also increase across the randomized 2, 4, and 6 mA conditions.

In those randomized active stimulation groups, hemodynamic responses referenced to the resting baseline showed progressively greater increases in HbT and HbO and decreases in HbR with increasing stimulation intensity, with the largest changes at 6 mA (Figure 3A.1-C.1). The intensity-dependent pattern persisted after correction using the time-matched 0 mA condition (Figure 3A.2-C.2). Across the 2, 4, and 6 mA, the mean within-participant dose-response slopes were 0.46 μM/mA for ΔHbT (95% CI, 0.29-0.63; P < 0.001), 0.56 μM/mA for ΔHbO (95% CI, 0.41-0.71; P < 0.001), and −0.10 μM/mA for ΔHbR (95% CI, −0.17 to −0.03; P = 0.010; Figure 3A.3-C.3). Sham was included as an additional control comparator but was not included in the active dose-response analysis.

**Figure 3.**
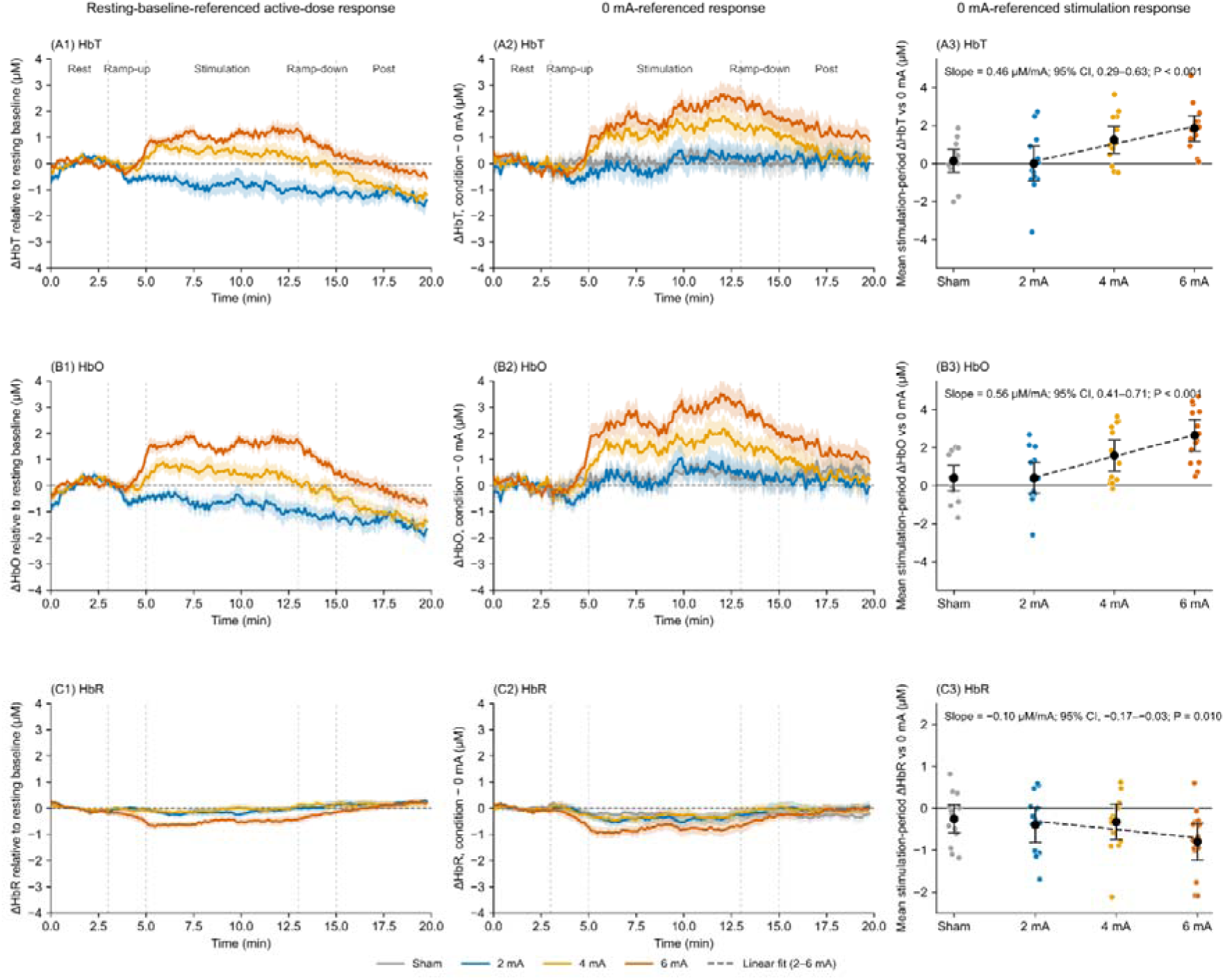
Intensity-dependent hemodynamic responses to HC-tDCS before and after 0 mA correction. (A.1–C.1) Baseline-referenced changes in HbT, HbO, and HbR, respectively, during active HC-tDCS at 2, 4, and 6 mA. Solid lines indicate group means, shaded bands indicate ±standard error of the mean (SEM), and gray vertical dashed lines mark stimulation phase boundaries. (A.2–C.2) Corresponding 0 mA-corrected responses under sham and active stimulation. For each participant, the 0 mA signal was subtracted pointwise from each condition, followed by referencing to the mean of the initial 3-min resting period. (A.3–C.3) Mean 0 mA-corrected ΔHbT, ΔHbO, and ΔHbR changes during the stimulation period. Colored points represent individual participants, and black circles with error bars indicate group means and 95% CIs. Dashed lines indicate mean within-participant dose-response slopes across 2, 4, and 6 mA; sham was included only as a control comparator.

Pairwise comparisons of the corrected responses showed greater ΔHbT and ΔHbO at 4 and 6 mA than under sham and at 2 mA (Figure 4). ΔHbO was also greater at 6 than at 4 mA, whereas the corresponding difference in ΔHbT did not remain significant after adjustment (P = 0.054). ΔHbR was lower at 6 mA than under sham and at 2 and 4 mA. No other pairwise differences were significant.

**Figure 4.**
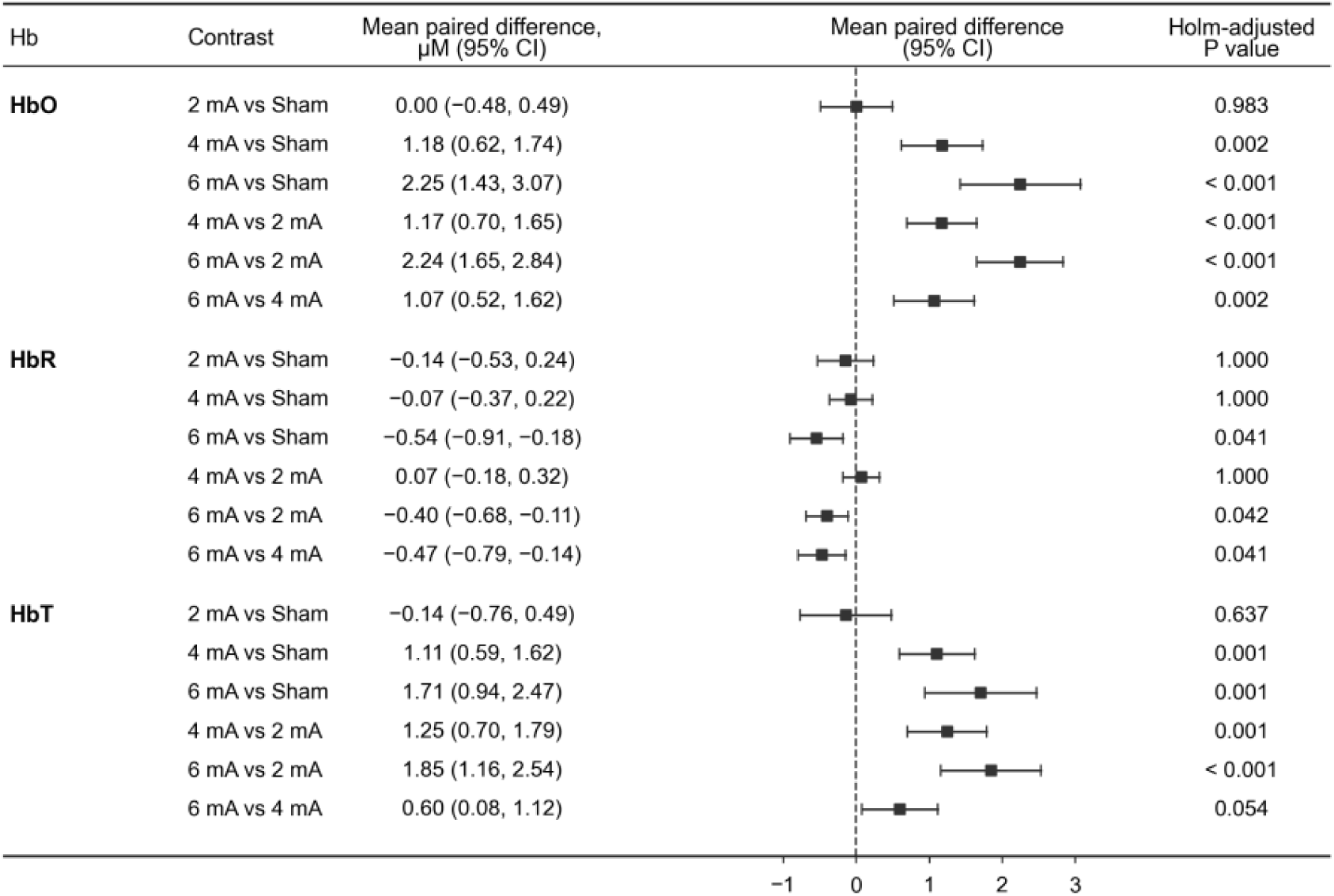
Pairwise comparisons of hemodynamic responses across HC-tDCS conditions. Pairwise differences in mean 0 mA-corrected stimulation-period ΔHbO, ΔHbR, and ΔHbT responses are shown for sham, 2 mA, 4 mA, and 6 mA conditions. Differences were calculated as the first-listed minus the second-listed condition and are presented as mean paired differences with 95% confidence intervals (CIs). P values were adjusted for six pairwise comparisons separately for each hemoglobin measure using the Holm procedure.

### Hemodynamic responses across source-detector distances

To assess whether the intensity-dependent hemodynamic responses extended to measurements with greater sensitivity to deeper tissue, we examined ΔHbT and ΔHbO across 4 source-detector separations (1.5, 2.12, 3.0, and 3.35 cm). Both measures were significantly greater than zero at all 4 separations during 4 and 6 mA stimulation. At 6 mA, mean stimulation-period ΔHbT values were 2.06, 1.81, 1.84, and 1.53 μM, respectively (all P < 0.01), while corresponding ΔHbO values were 3.24, 2.42, 2.53, and 2.00 μM (all P < 0.01; Figure 5).

**Figure 5.**
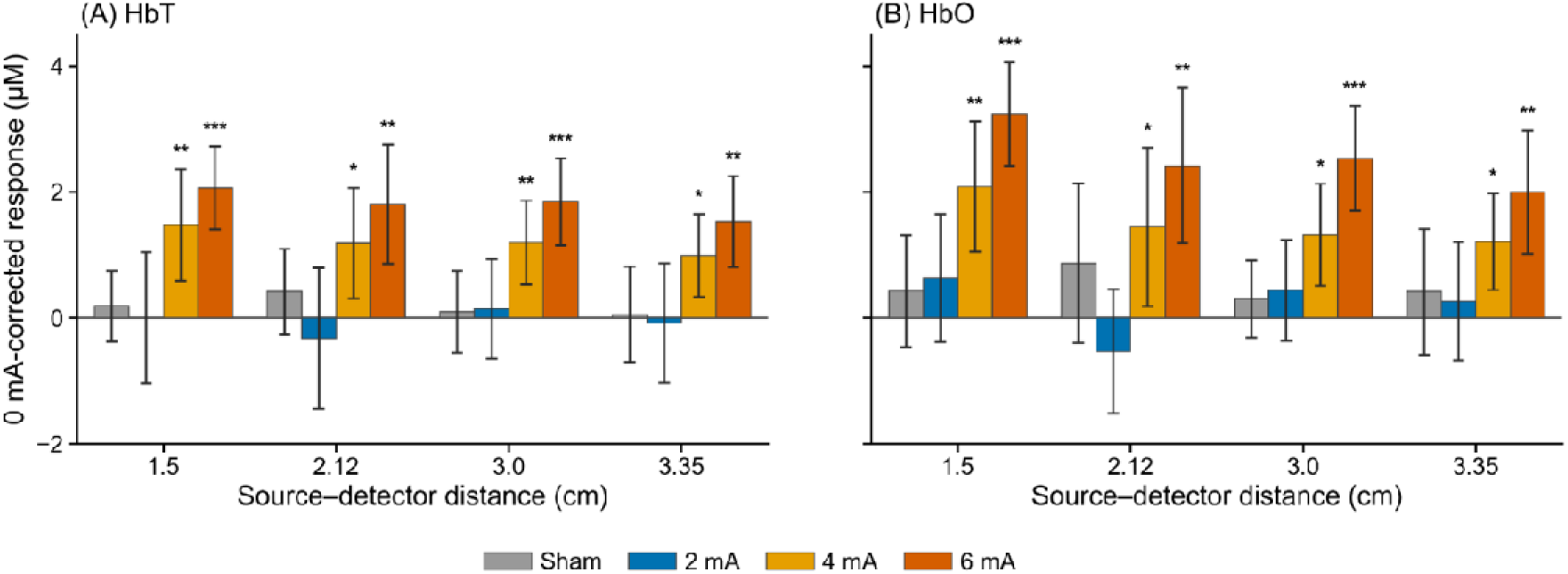
0 mA-corrected HbT and HbO responses across source–detector distances during HC-tDCS. (A) Mean stimulation-period ΔHbT under sham, 2, 4, and 6 mA conditions across source–detector separations of 1.5, 2.12, 3.0, and 3.35 cm. (B) Corresponding ΔHbO responses. Bars represent group means; error bars indicate 95% confidence intervals (CIs). For each condition and hemoglobin measure, participant-level responses at each separation were compared with zero using two-sided one-sample *t* tests. *P* values were Holm-adjusted across the four separations within each condition and hemoglobin measure. Asterisks indicate Holm-adjusted *P* values: \**P* < 0.05, \*\**P* < 0.01, and \*\*\**P* < 0.001.

Importantly, when analysis was restricted to the longer-separation channels (3.0 and 3.35 cm), both ΔHbT and ΔHbO remained significantly greater at 6 mA than under sham, 2 mA, and 4 mA stimulation. For ΔHbT, the mean paired differences were 1.66 μM versus sham (95% CI, 0.83-2.49; P = 0.004), 1.68 μM versus 2 mA (95% CI, 1.01-2.34; P < 0.001), and 0.61 μM versus 4 mA (95% CI, 0.20-1.02; P = 0.013). Corresponding differences for ΔHbO were 2.03 μM (95% CI, 1.08-2.97; P = 0.002), 1.98 μM (95% CI, 1.32-2.64; P < 0.001), and 1.06 μM (95% CI, 0.48-1.63; P = 0.006), respectively. For ΔHbR, only the difference between 6 and 4 mA remained significant (−0.45 μM; 95% CI, −0.74 to −0.15; P = 0.037; Figure S2).

### ΔHbT and ΔHbO slopes during stimulation period

To assess whether hemodynamic responses continued to develop during sustained stimulation, we quantified the temporal slopes of ΔHbT and ΔHbO over the 8-minute stimulation plateau. At 6 mA, both slopes were significantly positive: 0.19 μM/min for ΔHbT (95% CI, 0.08 to 0.30; P = 0.012) and 0.17 μM/min for ΔHbO (95% CI, 0.06 to 0.28; P = 0.021). Slopes under sham, 2 mA, and 4 mA did not differ significantly from zero (Figure 6). However, despite the significant positive slopes at 6 mA, the relationship between stimulation intensity and temporal slope across 2, 4, and 6 mA was not significant (ΔHbT: 0.02 μM/min/mA, 95% CI, −0.01 to 0.06, P = 0.146; ΔHbO: 0.02 μM/min/mA, 95% CI, −0.01 to 0.06, P = 0.217).

**Figure 6.**
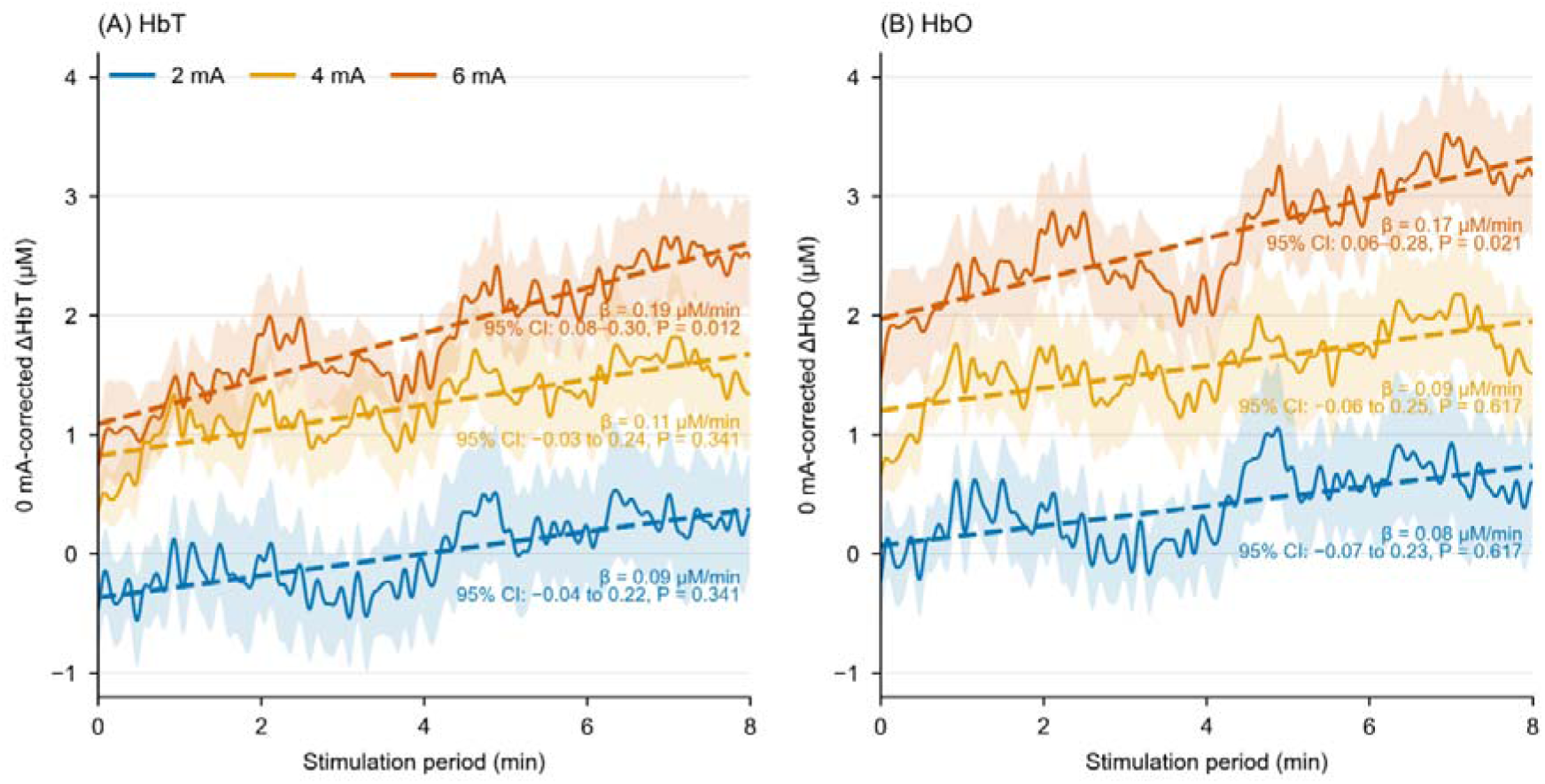
Hemodynamic response slopes during the stimulation period. (A) 0 mA-corrected ΔHbT responses during the 8-min stimulation period (5–13 min of the recording). (B) Corresponding 0 mA-corrected ΔHbO responses. Solid lines represent group mean responses, and shaded areas indicate ±SEM. Dashed lines show ordinary least-squares (OLS) fits to the group mean curves over the stimulation period. Annotations report the mean participant-level temporal slope (β, μM/min), 95% confidence interval (CI), and Holm-adjusted *P* value for each stimulation condition.

## Discussion

In this study, HC-tDCS produced a significant intensity-dependent hemodynamic response across 2-6 mA, characterized predominantly by increases in HbT and HbO, with the greatest responses at 6 mA. These responses were observed across all source-detector distances, including longer-separation channels. At 6 mA, HbT and HbO also continued to increase during the stimulation period. Together, these findings demonstrate a robust relationship between stimulation intensity and cerebral hemodynamic responses.

Previous studies of tDCS-induced hemodynamic responses have reported mixed findings [19]. Human perfusion studies have demonstrated changes in regional cerebral blood flow, but evidence for dose dependence has been inconsistent. Liu et al. found no consistent dose-dependent cerebral blood flow response during brief stimulation across 0.5-2 mA [15]. At higher intensities, Workman et al. reported substantial individual variability without a consistent group-level dose response across 1-4 mA [16], whereas Shinde et al. identified regional dose-dependent perfusion changes up to 4 mA [17]. Jamil et al. correlated imaged perfusion changes with current intensity and regional predicted electric field intensities [14]. Our findings extend these investigations to 6 mA and demonstrate a consistent increase in HbT and HbO across the tested intensities. Differences among studies may reflect measurement modality, stimulation recording period, reference condition / sham design, and electrode configuration.

In this last regard, the frontotemporal montage used for HC-tDCS produces ∼1.5-fold stronger brain electric fields per mA than conventional tDCS montages [6]. Consequently, the brain electric fields generated by 4 mA HC-tDCS would require ∼6 mA with these other montages, while those generated at 6 mA HC-tDCS are ∼2x and ∼4x those produced at 4 mA and 2 mA, respectively, in previous tDCS studies.

The observed hemodynamic pattern provides insight into the physiological effects of tDCS. Increasing stimulation intensity produced greater increases in HbT and HbO, accompanied by a smaller decrease in HbR. This pattern is compatible with greater delivery of oxygenated blood and a blood volume related hemodynamic response as stimulation intensity increases, rather than a response dominated by increased oxygen extraction. Such a hemodynamic phenotype could arise through direct vascular stimulation, neurovascular coupling, or both. Experimental studies support direct effects of electrical stimulation on vascular and perivascular structures [7]. Moreover, computational modeling by Arora et al. found that perturbation of the perivascular compartment best explained the early HbT response during tDCS [23]. Together, these observations support vascular stimulation as a potential contributor to the hemodynamic effects of tDCS, alongside neuronal and glial mechanisms.

The source-detector distance analysis provides further insight into the spatial distribution of these responses. Intensity-dependent increases in HbT and HbO were observed at all separations, and responses at 6 mA remained significantly greater than those at lower intensities when analysis was restricted to longer-separation channels (3.0 and 3.35 cm), which have greater sensitivity to deeper tissue [24]. A key limitation of fNIRS during tDCS is that stimulation may induce hemodynamic changes in both the scalp and brain. To address this, we used an active sham that maintained scalp stimulation while minimizing transcranial current flow. Results considering source-detector separation and active controls support a cerebral contribution to HC-tDCS induced hemodynamic changes.

Several limitations should be considered. The study was limited to healthy adults, and generalizability to clinical populations remains uncertain. The sham and 0 mA conditions were nonrandomized and conducted after the active sessions. Although longer-separation measurements and the active sham provided complementary information, the relative cerebral and superficial contributions to the hemodynamic responses could not be fully resolved.

### Conclusion

HC-tDCS produced intensity-dependent hemodynamic responses across 2-6 mA, with pronounced increases in HbT and HbO at 6 mA. These findings establish stimulation intensity as an important determinant of the hemodynamic response and support vascular stimulation as a potential contributor to the physiological effects of tDCS.

## Acknowledgments

We thank all participants for their time and contribution to this study.

## CRediT authorship contribution statement

Jinglin Mo: Conceptualization, Methodology, Investigation, Writing – original draft. Yahia Abdalla: Methodology, Software, Validation, Writing – review & editing. Mohamad FallahRad: Methodology, Resources, Writing – review & editing. Mojtaba Belali Koochesfahani: Methodology, Software, Writing – review & editing. Santiago Osorio Jurado: Methodology, Software, Writing – review & editing. Kyle Donnery: Methodology, Validation, Writing – review & editing. Marom Bikson: Conceptualization, Methodology, Supervision, Writing – review & editing.

## Ethics declaration

Written informed consent to take part in the study and to publish the article has been obtained from all participants or their legal representatives. The privacy rights of participants have been observed. This study was performed in compliance with relevant laws, regulatory frameworks and guidelines where the research took place. This study was approved by the City University of New York Institutional Review Board. (Approval No. 2024-0013-CCNY)

## Declaration of generative AI and AI-assisted technologies in the manuscript preparation process

During the preparation of this work, the authors used OpenAI Codex for language editing, journal-format adaptation, and manuscript organization. The authors reviewed and edited all AI-assisted content and took full responsibility for the content of the manuscript.

## Data availability

Data will be made available on request.

## Glossary

**fNIRS:** Functional near-infrared spectroscopy; **HC-tDCS:** High-capacity transcranial direct current stimulation; **HbO:** Oxygenated hemoglobin; **HbR:** Deoxygenated hemoglobin; **HbT:** Total hemoglobin; **OLS:** Ordinary least squares; **P99:** 99th-percentile electric-field magnitude; **SEM:** Standard error of the mean; **tDCS:** Transcranial direct current stimulation; **SD:** standard deviation.

